# Confounding effects of heart rate, breathing rate, and frontal fNIRS on interoception

**DOI:** 10.1101/2022.06.02.494474

**Authors:** Diego Candia-Rivera, M. Sofía Sappia, Jörn M. Horschig, Willy N. J. M. Colier, Gaetano Valenza

**Affiliations:** Bioengineering and Robotics Research Center E. Piaggio & Department of Information Engineering, School of Engineering, University of Pisa, 56122, Pisa, Italy; Artinis Medical Systems, B.V., Einsteinweg 17, 6662 PW, Elst, The Netherlands; Radboud University Nijmegen, Donders Institute for Brain, Behaviour and Cognition, 6525 EN, Nijmegen, The Netherlands

**Keywords:** fNIRS, heart rate variability, breathing control. heartbeat counting task, interoception

## Abstract

Recent studies have established that cardiac and respiratory phases can modulate perception and related neural dynamics. While heart rate and respiratory sinus arrhythmia possibly affect interoception biomarkers, such as heartbeat-evoked potentials, the relative changes in heart rate and cardiorespiratory dynamics in interoceptive processes have not yet been investigated. In this study, we investigated the variation in heart and breathing rates, as well as higher functional dynamics including cardiorespiratory correlation and frontal hemodynamics measured with fNIRS, during a heartbeat counting task. To further investigate the functional physiology linked to changes in vagal activity caused by specific breathing rates, we performed the heartbeat counting task together with a controlled breathing rate task. The results demonstrate that focusing on heartbeats decreases breathing and heart rates in comparison, which may be part of the physiological mechanisms related to “listening” to the heart, the focus of attention, and self-awareness. Focusing on heartbeats was also observed to increase frontal connectivity, supporting the role of frontal structures in the neural monitoring of visceral inputs. However, cardiorespiratory correlation is affected by both heartbeats counting and controlled breathing tasks. Based on these results, we concluded that variations in heart and breathing rates are confounding factors in the assessment of interoceptive abilities and relative fluctuations in breathing and heart rates should be considered to be a mode of covariate measurement of interoceptive processes.

## Introduction

In recent years, there has been a substantial increase in the number of studies on interoception that aim to describe the mechanisms involved in the sensing of inner bodily signals and their influence on brain dynamics.^1–3^ Interoception includes the sensing of cardiac, respiratory, and gut signaling.^4^ More in general, interoception has been defined as the sense of physiological condition of the entire body,^5^ which can potentially include other interoceptive sub-modalities, such as thermosensation, pain, or affective touch.^6^ Indeed, thermosensation, pain and affective touch involve a dynamic brain-heart interplay and associated interoceptive pathways.^7–9^ The functions associated with interoceptive processes include homeostatic and allostatic regulation as well as the integration of information that enables bodily awareness and consciousness.^1,3,10–13^ Evidence has demonstrated that neural responses to heartbeats contribute to visual,^14^ somatosensory,^15^ auditory,^16,17^ and self-perception.^18–20^ Further, recent evidence has suggested that breathing is aligned with the perception of sensory inputs,^21,22^ suggesting that it is involved in perceptual sensitivity modulation and, possibly, shaping neural dynamics.^23,24^

Heartbeat-evoked potential has been repeatedly reported as a metric for interoceptive awareness, for both comparing good heartbeat-sensing performances with bad ones and correlating it with the estimated interoceptive accuracy (early evidence provided by Pollatos and Schandry, 2004,^25^ a recent validation in heart-transplanted patients reported by Salamone et al., 2020,^26^ and a review by Coll et al., 2021 and Park and Blanke, 2019).^27,28^ Recent evidence has demonstrated that fluctuations in heart rate induce changes in stroke volume, which affects the estimation of heartbeat-evoked responses.^29^ These biases have been previously demonstrated in certain studies, revealing a relationship between the measured interoceptive accuracy and heart rate^30^ (for a review on biases, see Coll et al., 2021).^27^ Some studies have reported that observed effects in heartbeat-evoked potential are uncorrelated to changes in the mean heart rate and considered it as a control measure—however, to the best of our knowledge, no previous study has considered relative changes in heart rate or investigated cardiorespiratory variation during cardiac awareness tasks. Cardiorespiratory dynamics refer to the acknowledged influence of breathing on heart rate and blood pressure, and vice versa.^31^ This influence is most readily apparent in the case of the effect of breathing on heart rate—the increase of heart rate during inspiration and its decrease during expiration is termed respiratory sinus arrhythmia.^32^ Cardiorespiratory coupling has been acknowledged as a mutual, nonlinear, and bidirectional influence between breathing and heart rate.^31,33,34^

In this study, we comprehensively investigated variations in heart rate, cardiorespiratory correlation, and frontal hemodynamics during the performance of various tasks involving interoceptive communications. Moreover, we investigated the aforementioned physiological dynamics at paced breathing frequencies to distinguish between functional changes induced by interoceptive processes and those caused by mechanical breathing processes.^35^ However, our results did not conclusively establish whether or not cardiorespiratory dynamics are affected by the central nervous system when focusing on either the heartbeat or breathing or when the focus of the participant is on either monitoring or controlling visceral activity. We also explored variations in frontal hemodynamics, as measured using functional near-infrared spectroscopy (fNIRS), to verify the dependence of the effects on heart rate and cardiorespiratory correlation at different breathing rhythms on brain-body states.

We considered two tasks to perform neural monitoring and neural control of visceral activity. To measure the neural monitoring of visceral activity, the classic heartbeat counting task was used.^36^ To measure the neural control of visceral activity, a breathing control task based on Hernando et al., 2017 and Madhav et al., 2013 was used.^37,38^ The objectives of selecting these tasks were as follows. First, ascending communication from the viscera to the brain was stimulated by fostering cardiac awareness in the participant.^39,40^ Second, descending communication from the brain to the viscera was stimulated by controlling breathing rate.^41^

## Materials and Methods

### Participants

Twenty healthy young adults were recruited as participants in this study. Data from 19 subjects (age range: 21–34 years; median = 26 years; 8 males, 11 female) were used to study cardiorespiratory dynamics. A subset of 12 subjects (age range: 22–34 years; median = 26 years; 8 males, 4 females) was considered to study frontal hemodynamics, as measured using fNIRS (inclusion criteria are detailed below).

Of the 20 participants, one was excluded because of corrupted data and seven participants were not included in the fNIRS data analysis because of bad signal quality index.^42^

This study was approved by the local ethics committee Area Vasta Nord-Ovest Toscana. The subjects signed an informed consent form to participate in the study, all methods were performed in accordance with the relevant guidelines and regulations as required by the Declaration of Helsinki. Data acquisition was performed at Artinis Medical Systems B.V., Elst, The Netherlands. The instructions were provided in English language. None of the subjects declared any history of neurological, cardiovascular, or respiratory diseases.

### Experimental protocol

The recordings comprised fNIRS sampled at 50 Hz using a Brite23 device (Artinis Medical Systems B.V., the Netherlands, Fig. 1A), three-lead electrocardiography (ECG), and breathing inductive plethysmography (SleepSense, Elgin, USA). Both ECG and breathing signals were sampled at 250 Hz using a TMSi SAGA 32+/64+ amplifier (Twente Medical Systems International B.V., Netherlands). To record fNIRS signals, 23 channels with a source-detector separation of 3.5 cm were used to monitor brain activity in the frontal cortex (see Fig. 1A).

**Figure 1.**
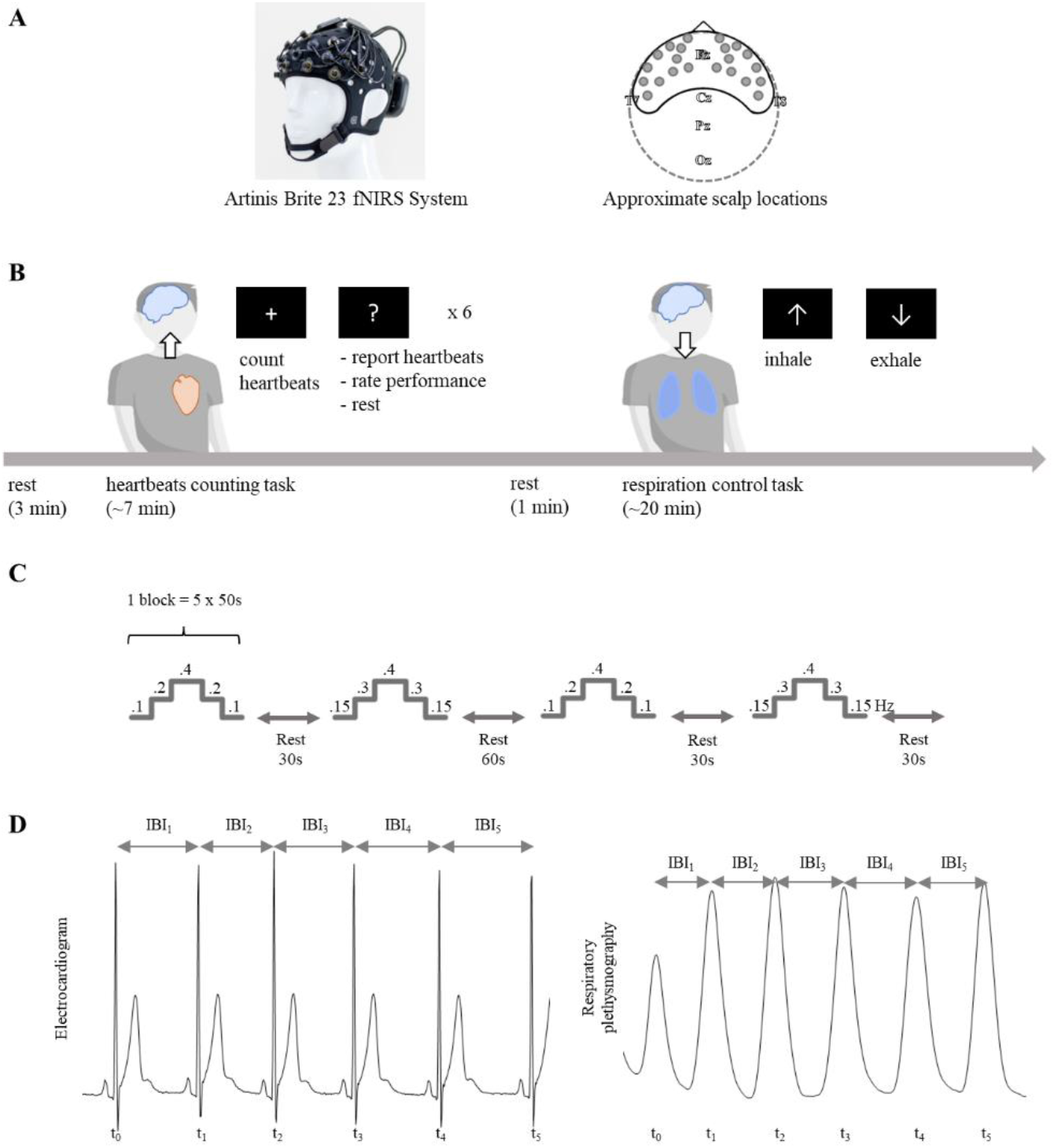
(A) Brite 23 (Artinis Medical Systems B.V., the Netherlands), the fNIRS system used in the experimental protocol. (B) Experimental protocol: heartbeat counting task and breathing control task. (C) Detailed transition during the breathing control task. (D) Estimation of heart and breathing rates.

The experimental protocol comprised a resting period lasting 3 minutes, a heartbeat counting task (7 minutes approximately), a resting period of 1 minute, and a breathing control task (20 minutes approximately), as depicted in Fig. 1B. Prior to the task, participants were asked to remove any accessories in their wrists and hands, such as watches, rings, or bracelets. Participants were located in a dark room and received visual instructions on screen (black background and white text).

#### Heartbeat counting task

Participants were asked to sit comfortably, put their arms on the table, and count their heartbeats by focusing on their corporeal sensations while fixating on a screen (without touching any body part with their fingers).^36^ The task was performed over six blocks of different durations between 30–60 seconds. The duration of each block was selected randomly for each trial and participant. No feedback was provided to the participants on their performance. After each block, the participants were explicitly asked to stop focusing on their heartbeats, report the number of heartbeats counted, and rate their self-perceived performance on a scale ranging from 1 (bad) to 9 (good).

The validity of some of these tasks has been questioned because of different biases.^30,43^ Alternatives have been proposed that reduce these biases.^44–46^ However, the objective of this study was to elicit cardiac awareness, rather than distinguish between the subjects based on their interoceptive abilities.^27^ To account for possible biases, trial durations was selected randomly to reduce the potential dependence of participants’ performance on trial duration and time perception. We explicitly asked participants to count only the heartbeats that they actually felt, rather than those that they expected to occur. As an additional control measure, we ensured that the participants’ interoceptive accuracy was related to their reported performance perception. The heartbeat counting task accuracy was computed based on the percentage error of the counted heartbeats with respect to the number of heartbeats detected via ECG, in accordance with the formula in Equation 1:

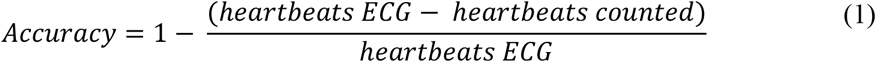

The computed markers associated with the heartbeat counting task were averaged over the six trials. In cases in which the transition between the performance and cessation of heartbeat counting was evaluated, the trials were aligned with the transitions. Statistical tests were performed 30 s before and after each transition to compare the respective autonomic- and hemodynamic-related markers.

After the task, participants were asked in which body part they mostly felt their heartbeats. The options were: head, chest, other body part, or in a diffuse way throughout the body. Participants were also asked whether they thought they estimated their heartbeats occurrences, rather than actually counting.

#### Breathing rate control task

The breathing control task was based on the works of Hernando et al. (2017) and Madhav et al. (2013).^37,38^ The participants were guided to inhale and exhale at a constant pace and at specific rates. The guidance consisted of a bar moving vertically together with a text indicating inhale or exhale phase. The protocol consisted of two blocks separated by a 30-second resting period, followed by a 60-second resting period. Subsequently, both blocks were repeated (see Fig. 1C). Each block comprised five steps in which the breathing rate was constant over a period of 50 s. The step transitions of the two blocks can be expressed as follows: 0.1 – 0.2 – 0.4 – 0.2 – 0.1 Hz, and 0.15 – 0.3 – 0.4 – 0.3 – 0.15 Hz.

Physiological data analysis was performed between transitions from slower to faster rates during the breathing control task: 0.1 – 0.2, 0.15 – 0.3, 0.2 – 0.4, and 0.3 – 0.4 Hz, and between transitions from faster to slower rates: 0.2 – 0.1, 0.3 – 0.15, 0.4 – 0.2, and 0.4 – 0.3 Hz.

During the entire protocol, each breathing rate occurred four times and each breathing rate transition occurred twice. The computed markers were averaged over all trials (i.e., individual rates were averaged over four trials and breathing rate transitions were evaluated by averaging over two trials).

### Data processing

Physiological and behavioral data were preprocessed using MATLAB R2018a and the Fieldtrip Toolbox.^47^

#### ECG processing

The ECG series were processed using an automated procedure and then visually inspected to verify their quality. The ECG were bandpass filtered with a Butterworth filter of order 4 between 0.5 and 45 Hz. On each ECG series, R-peak detection was performed using a peak template-matching method^48^ to derive heart rate variability series. All detected peaks were visually inspected over the original ECG series along with the inter-beat interval histogram. Ectopic detection was automatically performed by identifying the local maxima of the derivatives of the inter-beat interval series. Manual corrections through interpolation were performed as required.

The power spectrum density (PSDs) of heart rate variability series was computed using Burg’s method,^49^ with order 16 as recommended in Task Force of the European Society of Cardiology the North American Society of Pacing.^50^ The primary frequency of an inter-beat interval series was identified as the maximum peak in the 0–0.5 Hz interval of the PSD to characterize the dominant frequency.

#### Breathing rate signal processing

The breathing series were processed using an automated procedure and then visually inspected to verify their quality. The breathing rate series were smoothed using a 2-second-mean sliding time window based on inhalation peak detection and a peak template-matching method. All detected peaks were visually inspected over the original breathing series along with the inter-peak interval histogram. Ectopic detections were automatically identified by detecting the local maxima of the derivatives of the inter-beat intervals. Manual corrections were performed as required.

The computation of the time-varying heart and breathing rates was performed through the interbeat intervals (IBIs) and computed as shown in Equation 2 and Fig. 1D The ratio between heart and breathing rates was computed as well.^51^

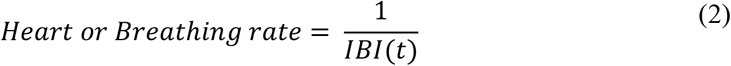

The breathing rate PSD were computed using Burg’s method of order 16. The primary frequency of an inter-beat interval series was identified as the maximum peak in the 0–0.5 Hz interval of the PSD.

#### Cardiorespiratory correlation

Cardiorespiratory correlation was measured using the Spearman cross-correlation coefficient between the breathing signals and the heart rate time series. The heart rate series were smoothed using a 2-second-mean sliding time window to remove high-frequency components (> 0.5 Hz). Cross-correlation was computed with respect to lags between -30 and +30 s. The maximum absolute value of the correlation coefficient within the lag interval was taken to be the cardiorespiratory correlation coefficient. The lag corresponding to the maximum correlation coefficient was recorded for further analysis.

#### fNIRS processing

Light intensities measured at 760 and 840 nm and digitized using 16-bit resolution were converted to optical densities, which, in turn, were converted into changes in concentration of oxygenated hemoglobin by applying the modified Beer-Lambert law.^52^ Oxygenation hemoglobin series were utilized because of their superior predictive performance compared to deoxygenation.^53^ The fNIRS data were bandpass filtered with a Butterworth filter of order 3 between 0.05 and 0.6 Hz to eliminate most of the systemic artifacts associated with pulses and very slow oscillations. Any systemic artifacts still remaining in the data were assumed to have minor impact on the results. The functional connectivity between channels was analyzed by comparing Spearman correlation coefficients.^54^ As only frontal hemodynamic activity was measured, connectivity analyses were performed for all possible combinations. For each channel, the median correlation coefficient computed with respect to all channels was considered.

#### Breathing response in fNIRS

We estimated the influence of breathing on fNIRS oxygenation signals, i.e., the breathing response and its variability under the studied conditions. The lag of the extracted breathing response corresponding to each signal was compared to the breathing signal recorded using the chest belt.

The breathing response was estimated in individual fNIRS channels.^55^ Because breathing components cannot be removed by filtering the signals,^56^ we separated them using principal component analysis (PCA). PCA was computed using 23 oxygenated hemoglobin channels and breathing signals. Individual components were analyzed by computing Spearman cross-correlation over a lag range of -30–30 s with respect to the breathing signal. Posterior component selection was performed based on the absolute maximum correlation coefficient over the aforementioned lag range. The resulting principal components with Spearman correlation coefficients higher than 0.6 were combined to reconstruct the breathing response from the fNIRS signals.

### Statistical analysis

Statistical analysis methods included the Spearman correlation coefficient, Wilcoxon’s signed-rank test for paired samples, and Friedman’s test for paired samples. All statistical tests on fNIRS markers were performed channel-wise.

Comparisons include rest vs. each experimental condition (heartbeats counting and breathing rate control). We also performed a comprehensive statistical analysis of the transitions between experimental conditions: counting heartbeats vs. not counting heartbeats, as well as transitions between breathing rates.

The *p*-values of the Spearman correlation coefficient were derived using a t-Student distribution approximation. The initial resting state was compared with the five rates of the breathing control task in the Friedman test. The significance levels of Spearman’s and Friedman’s *p*-values were corrected in accordance with the Bonferroni rule considering the number of fNIRS channels, with a corrected statistical significance of α = 0.05/23.

The Wilcoxon signed-rank test was performed using the normal approximation method. The significance levels of the *p*-values were corrected using a Monte Carlo permutation test with an initial critical α of 0.05. Samples were randomized over 10.000 permutations, and the Monte Carlo *p* was computed as the proportion of *p* with lower values. The significance of the Monte Carlo *p* was taken to be α = 0.05.

## Results

In this study we comprehensively investigated variations in heart rate, breathing rate, cardiorespiratory correlation, and frontal hemodynamics in fNIRS during the performance of various tasks involving interoceptive communications, as measured during heartbeats counting task and breathing rate control task.

In the heartbeat counting task, participants reported the body part in which they mostly felt their heartbeats. Three answered in the head, 8 in the chest, 4 in another body part, and 4 “in a diffuse way throughout the body”. Only 2 out of 19 participants reported that, at some point, they estimated their heartbeats occurrences rather than actually counting the heartbeats they felt.

### Average breathing and heart rates in each condition

We quantified the heart and breathing rates during the heartbeat counting and breathing rate control tasks. The heart rate and breathing rate group medians under all conditions are presented in Table I. Fig. 2 depicts the individual values of the heart and breathing rates. The heartbeat counting task did not cause significant modulation of the baseline heart and breathing rates. In contrast, the breathing control task induced a group-wise increase in heart rate with respect to the resting period corresponding to all the controlled breathing rates considered.

**Figure 2.**
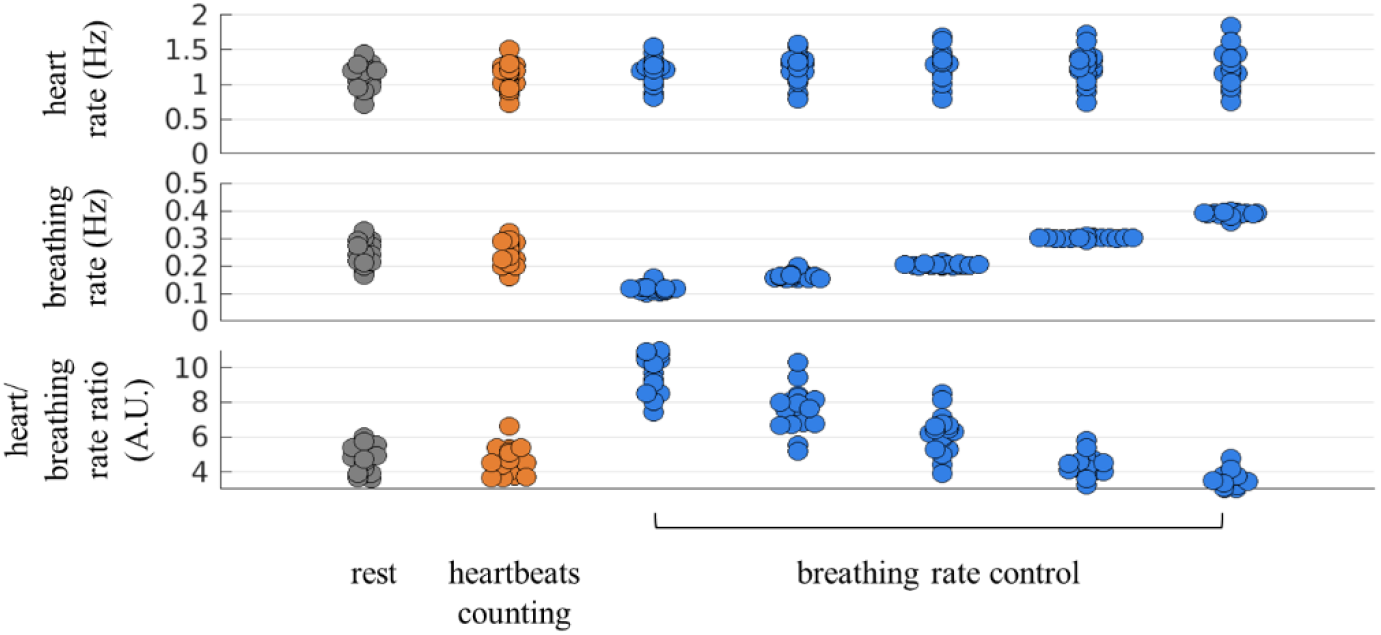
Average heart rate, breathing rate, and heart/breathing rate ratio during the experimental conditions.

**Table I.** Group median + MAD of the average heart and breathing rates in each experimental condition. P-values were computed using the paired Wilcoxon test during each session with respect the resting condition.

|  | Rest | Heart beat counting task | Test | Breathing control task |  |  |  |  |  |  |  |  |  |
| --- | --- | --- | --- | --- | --- | --- | --- | --- | --- | --- | --- | --- | --- |
|  |  |  |  | 0.1 | Test | 0.15 | Test | 0.2 | Test | 0.3 | Test | 0.4 | Test |
| Heart rate (Hz) | 1.18 ± 0.11 | 1.17 ± 0.11 | p=0.4939<br>Z=0.6841 | 1.21 ± 0.10 | **p=0.0015**<br>**Z=3.1791** | 1.27 ± 0.09 | **p=0.0003**<br>**Z=3.6217** | 1.29 ± 0.11 | **p=0.0002**<br>**Z=3.7828** | 1.27 ± 0.11 | **p=0.0003**<br>**Z=3.6218** | 1.30 ± 0.13 | **p=0.0002**<br>**Z=3.7425** |
| Breathing rate (Hz) | 0.24 ± 0.03 | 0.25 ± 0.04 | p=0.6292<br>Z=0.4829 | 0.11 ± 0.00 | - | 0.16 ± 0.00 | - | 0.20 ± 0.00 | - | 0.30 ± 0.00 | - | 0.39 ± 0.00 | - |
| Heart /Breathing rate ratio (A.U.) | 4.83 ± 0.69 | 4.54 ± 0.55 | p=0.6009<br>Z=-0.5231 | 10.48 ± 1.10 | **p=0.0001**<br>**Z=3.823** | 7.75 ± 0.57 | **p=0.0001**<br>**Z=3.823** | 6.33 ± 0.43 | **p=0.0002**<br>**Z=3.6218** | 4.22 ± 0.34 | **p=0.0329**<br>**Z=-2.1328** | 3.34 ± 0.34 | **p=0.0002**<br>**Z=-3.7828** |

### Relative changes in breathing and heart rates during transitions

Subsequently, we quantified the relative changes in heart and breathing rates. Each comparison during the heartbeat counting task was performed with respect to the subsequent resting period. In the breathing control task, the participants transitioned from slower to faster and from faster to slower breathing rates. The breathing and heart rate series were aligned during each transition, and z-score normalization was applied over the interval, -30–30 s, in the heartbeat counting task, and -50–50 s in the breathing control task.

As depicted in Fig. 3A, the transition between the performance and cessation of the heartbeat counting task induced a relative increase in the heart and breathing rates. Table II presents the group medians of these relative increases. The increase in heart rate was significant, whereas that in breathing rate was not significant corresponding to time windows of -30–30 s with respect to the transition. The results also revealed that the increase in breathing rate preceded the increase in heart rate, and that the decrease in the breathing rate was rapid whereas the higher heart rate was sustained for approximately 15 s. Table II also displays the change in the predominant frequency in the PSD, which was also significant only for the heart rate.

**Figure 3.**
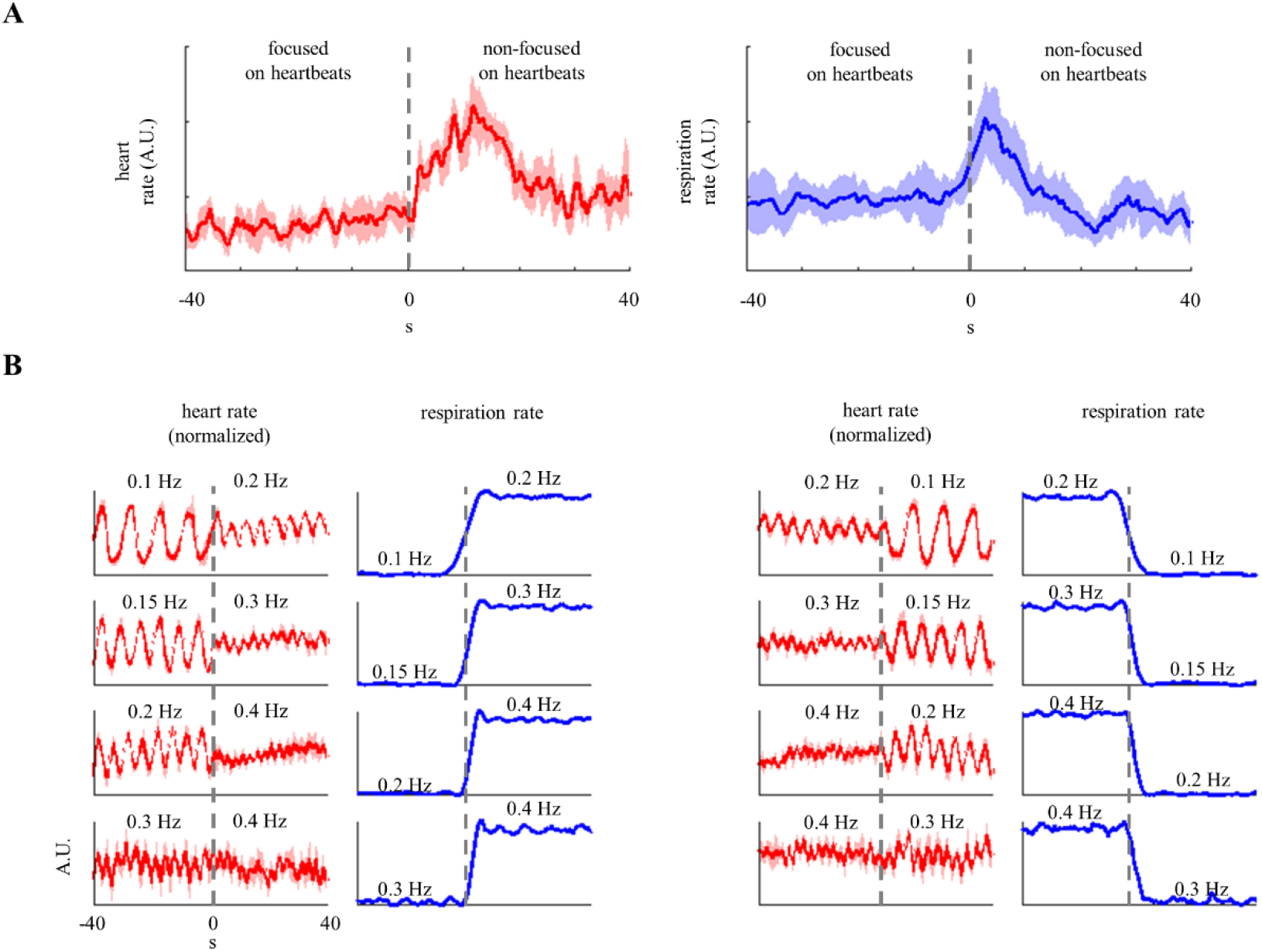
Respiratory modulations induced on heart rate. (A) Group median + MAD representing relative change in heart and breathing rates, triggered by the transition from the performance to the cessation of focusing on heartbeats. Heart rate data were smoothed using a 10-second moving mean window and a sample step of 1. Each subject data corresponded to an average over six trials. (B) Relative change in heart (group median + MAD) and breathing rates during slower-to-faster and faster-to-slower transitions. Each subject data corresponded to the average over the two trials for each transition. All signals represent the group median ± median absolute deviation. All signals used for computing the medians were the rates expressed in Hz, the z-score was normalized per subject over a window of -50–50 seconds with respect to each transition in breathing rate.

**Table II.** Group median + MAD of the average normalized heart and breathing rates and the main frequency in the PSD of the heart and breathing rates during the heartbeats counting task. P-values were computed using the paired Wilcoxon test over a 30-second period of counting heartbeats and a subsequent 30-second resting period.

|  | Mean rate (normalized, A.U.) |  |  |  | Main frequency (Hz) in the PSD |  |  |  |
| --- | --- | --- | --- | --- | --- | --- | --- | --- |
|  | Counting heartbeats | Not counting heartbeats | p | Z | Counting heartbeats | Not counting heartbeats | p | Z |
| Heart rate | -0.35 ± 0.13 | 0.35 ± 0.13 | **0.0001** | **3.823** | 0.12 ± 0.05 | 0.04 ± 0.04 | **0.0019** | **-3.0986** |
| Breathing rate | -0.02 ± 0.18 | 0.02 ± 0.18 | 0.2954 | 1.0463 | 0.07 ± 0.01 | 0.07 ± 0.02 | 0.2388 | -1.178 |

Fig. 3B depicts the relative changes while switching from one breathing rate to another. A consistent change in heart rate oscillations was observed—a fixed breathing rate triggered relative heart rate oscillations at the same breathing frequency. The modulation from breathing to heart rate oscillations was observed independently with respect to the changes corresponding to slower-to-faster or faster-to-slower breathing rates. Table III records the main group-median frequencies of the heart rate variability PSD. The main frequencies matched only the breathing rate corresponding to the 0.1-0.2 and 0.2-0.1 Hz transitions. Corresponding to breathing rates of 0.3 and 0.4 Hz, the predominant frequency remained ∼0.1 Hz. These results establish that fast breathing rates (> 0.2 Hz) triggered heart rate oscillations at the same breathing frequency. However, the main frequency remained in the low-frequency range.

**Table III.** Group median + MAD of the main frequency obtained from heart rate variability PSD during the breathing control task. P-values were derived using the paired Wilcoxon test during each session with respect the resting period.

| Slower-to-faster transition in breathing rate (Hz) | Main frequency (Hz) obtained from heart rate variability PSD |  |  |  | Faster-to-slower transition in breathing rate (Hz) | Main frequency (Hz) obtained from heart rate variability PSD |  |  |  |
| --- | --- | --- | --- | --- | --- | --- | --- | --- | --- |
|  | at slower breathing rate | at faster breathing rate | p | Z |  | at faster breathing rate | at slower breathing rate | p | Z |
| 0.1→0.2 | 0.098 ± 0.003 | 0.195 ± 0.004 | **0.0045** | **2.805** | 0.2→0.1 | 0.195 ± 0.008 | 0.098 ± 0.004 | **0.0008** | **-3.3513** |
| 0.15→0.3 | 0.148 ± 0.004 | 0.094 ± 0.012 | 0.3340 | -0.9661 | 0.3→0.15 | 0.086 ± 0.023 | 0.145 ± 0.004 | 0.3543 | 0.9263 |
| 0.2→0.4 | 0.199 ± 0.004 | 0.089 ± 0.012 | 0.4078 | -0.8278 | 0.4→0.2 | 0.086 ± 0.043 | 0.195 ± 0.008 | 0.7626 | 0.302 |
| 0.3→0.4 | 0.086 ± 0.023 | 0.094 ± 0.089 | 0.6008 | 0.5233 | 0.4→0.3 | 0.098 ± 0.094 | 0.098 ± 0.016 | 0.6012 | -0.5227 |

### Cardiorespiratory correlation

Cardiorespiratory correlation was measured in terms of the correlation coefficient between breathing and heart rate series at frequencies < 0.5 Hz. The median correlation at rest was 0.26 (on a scale ranging from 0 to 1). The heartbeat counting task induced a significant increase in the correlation, increasing the median to 0.47. During the breathing control task, the cardiorespiratory correlation was higher than that during the resting period only corresponding to the transitions between 0.2 and 0.4 Hz and 0.3 and 0.4 Hz (see Table IV). These results indicated that, under paced and controlled breathing, cardiorespiratory correlation was higher when the breathing rate was in the high-frequency band.

**Table IV.** Group median + MAD of cardiorespiratory correlation in each experimental condition. P-values correspond to the paired Wilcoxon test of each session with respect the resting condition.

|  | Rest | Heartbe at counting task | Test | Breathing control task |  |  |  |  |  |  |  |
| --- | --- | --- | --- | --- | --- | --- | --- | --- | --- | --- | --- |
|  |  |  |  | 0.1→0.2 Hz | Test | 0.15→0.3 Hz | Test | 0.2→0.4 Hz | Test | 0.3→0.4 Hz | Test |
| Cardioresp iratory correlation | 0.26 ± 0.01 | 0.47 ± 0.08 | **p=0.0006**<br>**Z=3.4206** | 0.29 ± 0.06 | p=0.6292<br>Z=0.4829 | 0.37 ± 0.06 | p=0.099<br>Z=1.6499 | 0.42 ± 0.13 | **p=0.0298**<br>**Z=2.1731** | 0.37 ± 0.04 | **p=0.0298**<br>**Z=2.1731** |
|  |  |  |  | 0.2→0.1 Hz | Test | 0.3→0.15 Hz | Test | 0.4→0.2 Hz | Test | 0.4→0.3 Hz | Test |
|  |  |  |  | 0.28 ± 0.09 | p=0.6874<br>Z=-0.4024 | 0.31 ± 0.05 | p=0.4445<br>Z=0.7646 | 0.44 ± 0.10 | **p=0.0112**<br>**Z=2.5353** | 0.28 ± 0.05 | **p=0.0269**<br>**Z=2.2133** |

We further explored the relationships triggered during the heartbeat counting task. First, we quantified the lag in the maximum correlation coefficient between the heart and breathing rates, as depicted in Fig. 3A. We computed the Spearman correlation coefficient between the two signals, considering a lag range of -20–20 s. As depicted in Fig. 4A, maximum correlation was observed corresponding to a lag of approximately 10 s. Further, as a control measure, we investigated the possible dependence of cardiorespiratory correlation on different interoceptive dimensions. No significant correlation was observed between cardiorespiratory correlation and the estimated interoceptive accuracy (Spearman R = -0.1123, p = 0.6465, Fig. 4B). Cardiorespiratory function was also observed to be uncorrelated to the participants’ perceived performance or self-confidence accuracy, which was reported without prior feedback (Spearman R = 0.1186, p = 0.6287, Fig. 4 C). As a control measure, we verified that the interoceptive accuracy is related to the performance, rather than being an estimation of each trial’s duration by confirming a significant correlation between the interoceptive accuracy and self-confidence scores (Spearman R = 0.6632, p = 0.0020, Fig. 4D).

**Figure 4.**
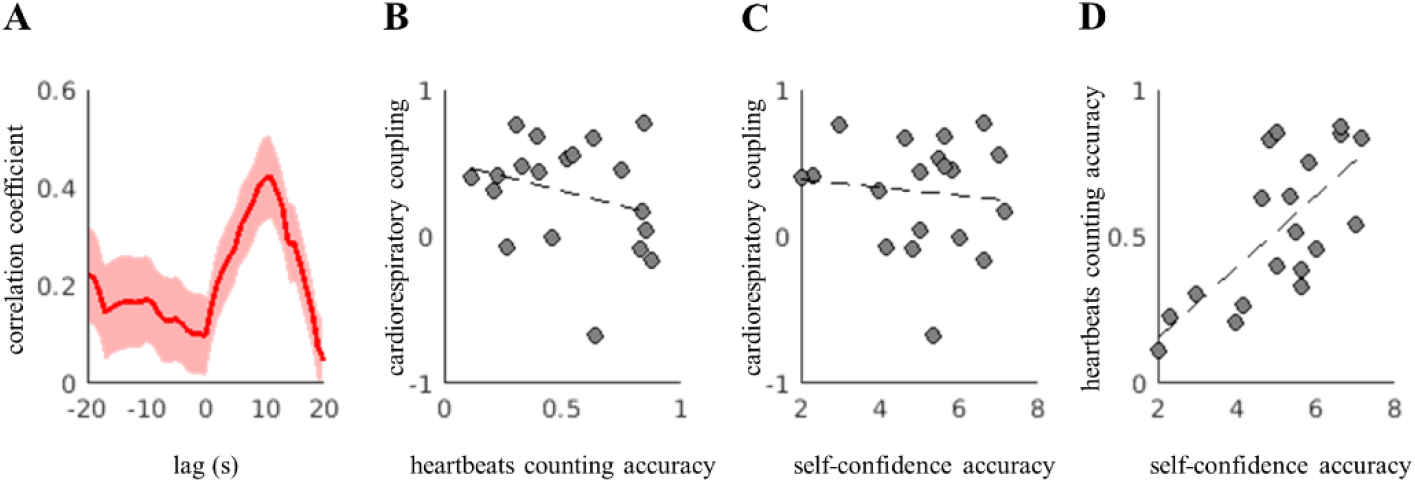
(A) Cardiorespiratory correlation lag, and the relationships between (B) cardiorespiratory coupling and heartbeat counting accuracy, (C) cardiorespiratory correlation and self-confidence regarding heartbeat counting, and (D) heartbeat counting accuracy and self-confidence regarding heartbeat detection.

### Frontal hemodynamics

We quantified changes in frontal hemodynamics during the performance of the two aforementioned tasks. First, during the heartbeat counting task, hemodynamics was estimated by comparing the changes in connectivity between different segments of heartbeat counting with respect to consecutive segments of rest. We observed increased frontal connectivity when participants focused on their heartbeats compared to when they did not. Changes in frontal connectivity were observed in the bilateral frontal channels but not in the midline channels, as illustrated in Fig. 5A. We further explored variations in connectivity during transitions in breathing rate. As illustrated in Fig. 5B a decrease in connectivity was observed during the transition from a slower to a faster breathing rate, and an increase in connectivity was observed during the transition from a faster to a slower breathing rate. Further, the scale of the effect was lower corresponding to transitions from faster breathing rates, whether increasing or decreasing. We then compared the differences in respiratory response latency in the fNIRS with respect to the five different breathing rates studied. To this end, a Friedman test was performed for each channel to compare the latencies of the respiratory responses under all conditions. As depicted in Fig. 5C, the major changes in the respiratory response with respect to the different breathing rates were observed in the midline channels, as well as in the far bilateral channels. As illustrative examples, Fig. 5D presents fNIRS signals for one trial of heartbeat counting task and breathing control task in the cases of increasing and decreasing breathing rate (between 0.1–0.2 Hz).

**Figure 5.**
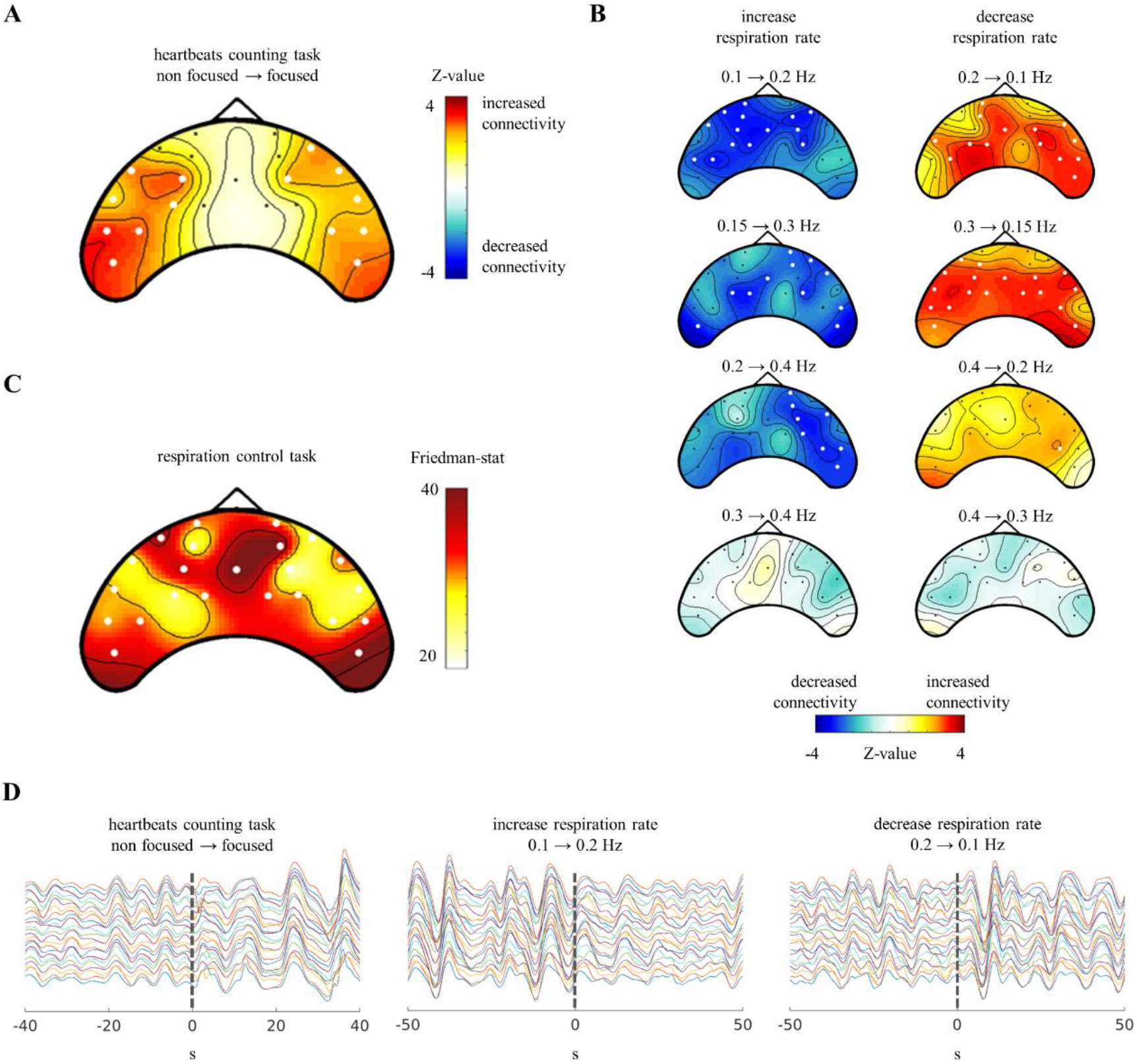
Frontal hemodynamics results. (A) Changes in connectivity induced by the cessation of focusing on heartbeats. (B) Changes in connectivity induced by switching the breathing rate in the breathing rate control task. (C) Changes in the respiratory response in fNIRS with respect to changes in breathing rhythm. Thick white channels indicate significance corrected for multiple comparisons. (D) Exemplary fNIRS signals for one trial of the heartbeats counting and breathing control tasks.

## Discussion

To highlight the acknowledged role of cardiac and breathing phases in different perceptual paradigms, in this study, we tested the role of heart rate variability and cardiorespiratory correlations during interoceptive processes such as heartbeat monitoring and guided breathing rate control. We described the heart rate, breathing rate, cardiorespiratory correlations, and frontal hemodynamics during the performance of these tasks.

### Heartbeats and breathing

The periods of heartbeat counting induced a relative decrease in breathing rate, followed by a relative decrease in heart rate. This indicated that the cessation of focusing on heartbeats induced a relative increase in the autonomic tone. Possible changes in the heart rate were controlled in previous studies, with non-significant changes in heart rate during heartbeat counting,^57^ as recorded in Table I and Fig. 2. However, in this study, after normalizing the trials for each subject, the relative change in the heart rate was observed to be persistent among the subjects, as depicted in Fig. 3A and Supplementary Figures 1 and 2. The relative changes in heart rate induced in our study was not voluntary. Indeed, heart rate cannot be voluntarily controlled through interoceptive awareness tasks,^58^ differently from voluntary heart rate modulations induced through neurofeedback techniques in specific conditions.^59^

Our results confirmed the presence of cardiorespiratory correlations under the two conditions studied—heartbeat counting and breathing rate control. Breathing wielded a clear influence on heart rate, instead of the other way around. Our results were in agreement with the repeated reports of the effect of breathing on heart rate.^60–64^ The influence of heart rate on breathing under the conditions studied was unclear, but if present, these influences exhibited significantly low magnitude.

Mechanisms that mediate the influence of breathing on cardiovascular function include autonomic afferents and efferents,^31,65,66^ as well as mechanical interactions.^35^ These effects are controlled in the brainstem—specifically in the pons.^67^ However, baroreceptors are hypothesized to mediate the mechanisms on the influence of the cardiovascular system on breathing.^31,68,69^ In this study, the influence of the breathing rate on heart rate exhibited a group-wise latency of 10 s during the heartbeat counting task. Previous studies that reported such delayed effects on heart rate also reported, for instance, a 16-second delay when holding breathing.^55^ These differences may be related to respiratory sinus arrhythmia and the physiological differences associated with the holding of breath and slow-paced breathing. However, such delays were not observed during the breathing control task in this study. Instead, the observed variations in heart rate variability during the breathing control task were instantaneous and associated with heart rate oscillations, rather than a specific increase or decrease in heart rate.

Previous studies have reported that delayed responses to breathing vary when the breathing rate is controlled with respect to normally paced breathing.^70^ The effects on heart rate variability oscillations have previously been described corresponding to paced breathing at ∼0.1 Hz.^71^ However, we observed heart rate oscillations corresponding to all breathing rates (Fig. 3B).

### Monitoring of visceral activity

We observed increased connectivity in frontal hemodynamics when subjects focused on their heartbeats. This increased connectivity was observed in the frontal and bilateral channels, but not in the midline channels. Instead, the breathing response exhibited higher covariation with respect to the breathing rate in the midline channels. As previously reported, resting state fluctuations in breathing rate covary with the default mode network midline frontal node.^72^ The midbrain periaqueductal gray also participates as an integrative structure in breathing interoception and has extensive projections to the ventrolateral structures.^73^ Our results reinforced the existing evidence that frontal hemodynamics are functionally involved in the monitoring of visceral activity, with some dissociation of heartbeats and breathing.

Previous studies have established that cortical responses to heartbeats are related to attention—both interoceptive or exteroceptive.^57^ Breathing has also been reported to modulate brain activity and map to canonical resting state networks, suggesting that breathing processing is irrespective of the conditions involved.^74^ The dissociation of interoception from cardiac and breathing signals also appears to be related to anxiety.^75,76^

The alignment of breathing with sensory perception^21,22^ is similar to cardiac phase alignments with somatosensory,^15^ auditory,^16,17^ and visual perception.^14^ The impact of breathing rate on cognitive functioning has already been demonstrated in mice and other animals. These relationships between breathing and cognitive processes suggest a specific role for breathing as a timing signal for brain activity.^77^ For instance, it has been shown that the breathing rate shapes olfactory memory,^78,79^ emotions,^80^ and reaction time,^81^ and coordinates the coupling of brain oscillations during sleep^82–84^ as well as during wakefulness.^85–90^ These results indicate that cardiorespiratory activity contribute to the shaping of attention and perception, and relative changes in breathing and heart rates are part of ongoing bidirectional brain-heart interplay processes,^91^ similar to emotions or stress.^7,92^

### Voluntary breathing control

We observed that frontal hemodynamics were actively involved during the breathing control task, either increasing or decreasing the breathing rate. Stronger effects were observed in slower transitions of breathing rates than in faster ones. These results indicate that frontal hemodynamics are involved in breathing control as well, as well as in non-uniform intensification of the presence of systemic components in the fNIRS signals. However, voluntary changes in breathing have been reported in the primary sensory and motor cortices, supplementary motor area, cerebellum, thalamus, caudate nucleus, and globus pallidus in the medulla,^41^ not in frontal activity. Frontal activations have been known to involve other processes, such as the focus of attention on breathing, which has been shown to activate the insula and deactivate the default mode network.^93^ Non-voluntary changes in breathing have different cerebral origins, including the default mode network midline frontal node,^72^ brain stem,^94^ and cerebellar vermis.^95^ Changes in frontal connectivity may also be related to the effort of breathing, given that guided breathing over long periods can cause discomfort due to muscle fatigue and suboptimal unloading of CO_2_ ^96^ and that the fatigue system involves the frontal cortex.^97^

### Clinical relevance

Our results indicated the presence of cardiorespiratory coupling under all conditions studied, with disruption. Disruptions in cardiorespiratory coupling are attributed to possible pathological conditions^98^, as well as affective states.^99–101^ Disrupted interoceptive abilities have also been associated to pathological conditions.^2,102–104^ However, the relationships between interoceptive abilities and mental health remain under debate.^105^ Further evidence exists on the relevance of cardiorespiratory communication with the central nervous system—for instance, the reported relationship between disrupted interoception abilities related to cardiac and breathing signals and anxiety.^75,76^ In addition, relationships with other visceral organs have been demonstrated by the apparent effects of gut microbiota on cardiorespiratory homeostasis.^106^

We found a significant correlation between the heartbeat counting task accuracy and the self-perceived performance. This correlation has been reported in other studies in healthy participants.^107,108^ Therefore, a loss of correlation, i.e., a mismatch between the obtained accuracy and self-perceived performance, may potentially relate to pathological conditions, such as neurodegeneration^103^ A better understanding of the physiological subtracts of a disrupted interoceptive awareness may lead to future developments with clinical relevance, given the abundant evidence linking interoception with mental health.^2,44^

### Limitations and future directions

Future research investigating interoceptive abilities should consider the registration of multiple peripheral signals to control the confounding factors. In this study was revealed that heartbeat counting task elicits a slower breathing rate and, consequently, slower heart rate. In light of the existing evidence about the effects of heart rate on heartbeat-evoked potentials,^29^ the change in breathing rate should be considered as confounding factor as well. As the present study was limited to fNIRS only, it remains to further explore how these changes in frontal hemodynamics relate to cortical electrophysiology, together with the potential influence of the change in cardiorespiratory correlation on heartbeat-evoked potentials.

A limitation of this study is the small sample size. Note that the group-wise effects may also be observed at a single-participant level (see Supplementary Material, Fig. 1 and 2).

The modifications to the heartbeat counting task, such as the explicit request to only report the heartbeats felt, contribute to reduce potential biases, including the time estimation.^109^ In this study, the instruction to not estimate the heartbeats occurrences and exclusively count the heartbeats felt was effective in 17 out 19 participants, according to the self-reports. It is worth mentioning that other interoceptive modalities have recently been explored, including breathing,^110^ while other interoceptive sub-modalities are still under debate.^6^ The heart rate has been reported as a confounding factor for measurements of interoceptive accuracy^27^ and heartbeat-evoked potentials.^29^ However, heart rate changes seem not correlated with the participants’ performance on counting or detecting their heartbeats.^44^ These differences have also been hypothesized to relate with individual differences on prior bodily representations and prediction errors.^111^ Given that attention has the effect of optimizing precision, within and between sensory modalities, individual differences may relate to the level of prioritization of interoception over other sensory modalities.^111^ These differences may also explain the enhanced interoceptive abilities reported in blind individuals.^112^

Because of its modulation effect, we emphasize the relevance of analyzing respiratory dynamics in cardiac awareness tasks. Indeed, participants are more likely to detect heartbeats during inhalation, as compared to exhalation.^113^ In light of the previously reported neuroimaging correlates of interoception,^40^ we also emphasize the breathing rate modulation effect on frontal hemodynamics.

## Conclusions

This study contributes to the understanding of physiological processes sustaining cardiorespiratory coupling that are relevant to the study of pathological conditions that disrupt both coupling and interoceptive abilities. Our results indicated that heart rate, breathing rate, and cardiorespiratory dynamics may act as confounding factors in the assessment of interoceptive abilities. Further analysis of cardiorespiratory coupling and associated brain dynamics is expected to reveal additional details regarding brain viscera functioning during the performance of cardiac awareness tasks.

## Author contributions

D.C.-R. conceived the study. D.C.-R., M.S.S., and J.M.H. designed the study. D.C.-R., and M. S.S. acquired the data. D.C.-R. Processed the data and figures. D.C.-R., M.S.S., and G. V. analyzed the data. D.C.-R., M.S.S. and G. V. wrote the manuscript. All authors (D.C.-R., M.S.S., J.M.H., W.N.J.M.C. and G. V.) have revised and approved the manuscript.

## Acknowledgements

The authors thank Naser Hakimi and Aline Simonetti for their help in acquiring the data.

## Data availability statement

Data are available upon request to the corresponding author.

