## Supplementary material for "Confounding effects of heart rate, breathing rate, and frontal fNIRS on interoception"

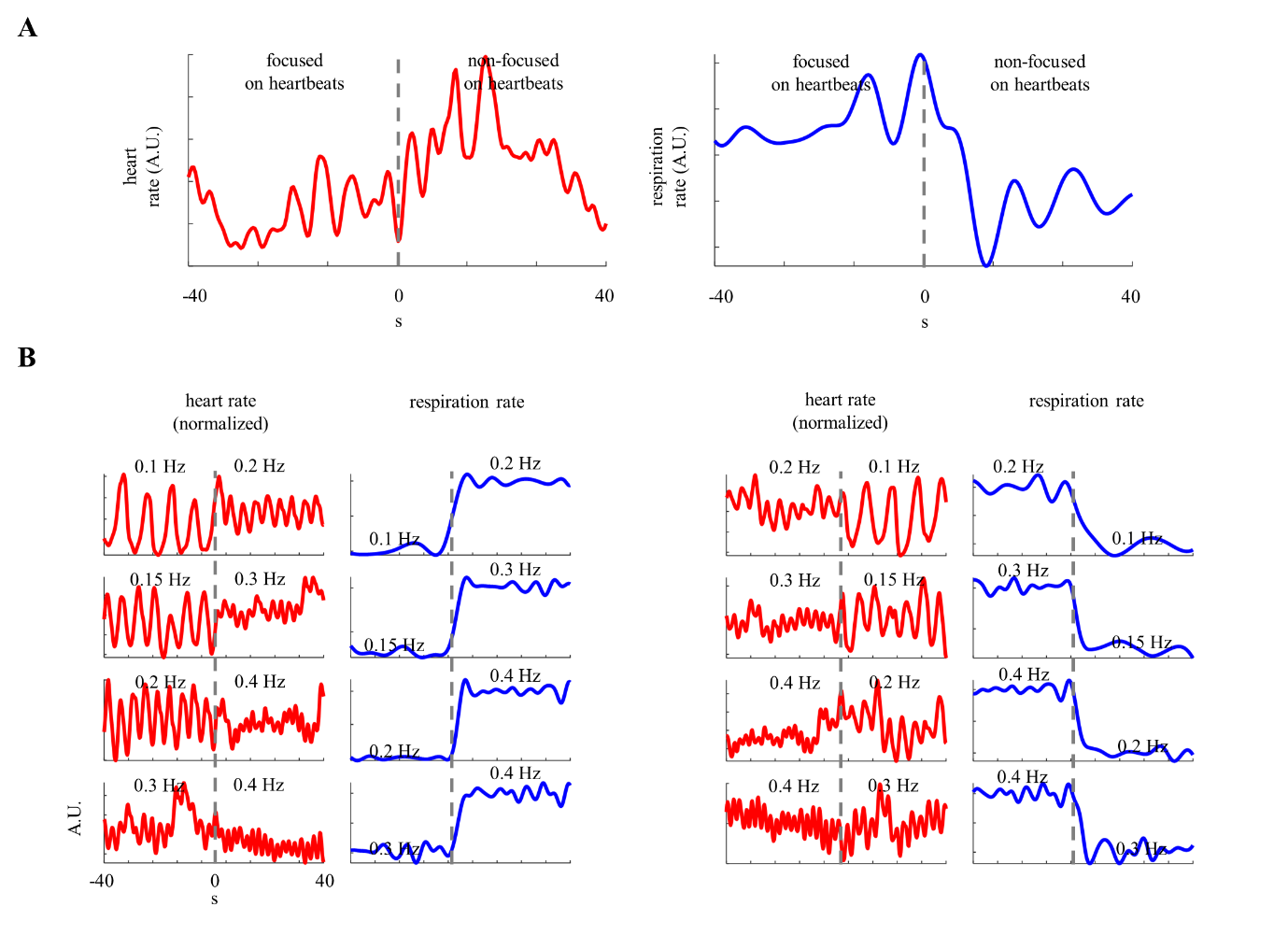

Supplementary Figure 1. Induced respiratory modulations on heart rate in a single subject. (Panel 1 A): Relative change in heart and breathing rates triggered by the transition from the performance to the cessation of focusing on heartbeats. Heart rate series were smoothed using a 10-second moving average window and a one-sample step. Data corresponds to an average over six trials. (Panel B) Relative change in heart and breathing rates expressed in Hz during slower-to-faster and faster-to-slower breathing rate transitions. All signals are z-score normalized over a window of -40–40 seconds with respect to each transition in breathing rate.

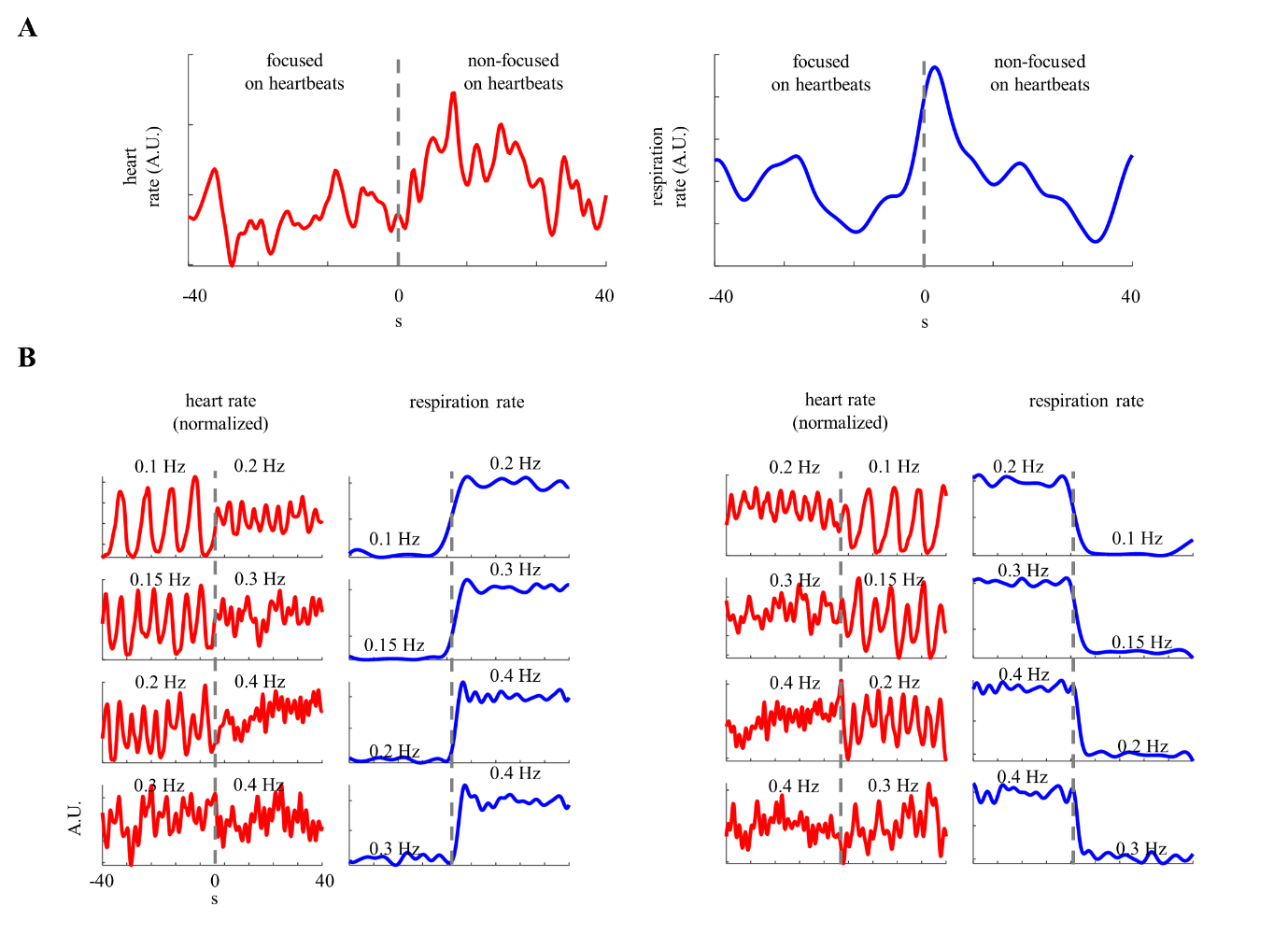

Supplementary Figure 2. Induced respiratory modulations on heart rate in a single subject. (Panel 1 A): Relative change in heart and breathing rates triggered by the transition from the performance to the cessation of focusing on heartbeats. Heart rate series were smoothed using a 10-second moving average window and a one-sample step. Data corresponds to an average over six trials. (Panel B) Relative change in heart and breathing rates expressed in Hz during slower-to-faster and faster-to-slower breathing rate transitions. All signals are z-score normalized over a window of -40–40 seconds with respect to each transition in breathing rate.
